# Climatic niches provide insights into the evolutionary origins and ecological significance of the succulent CAM syndrome around the world

**DOI:** 10.1101/2023.05.02.539181

**Authors:** Tania Hernández-Hernández, Marilyn Vásquez-Cruz, Israel Loera, Melina DelAngel, Miguel Nakamura

## Abstract

Although distributed globally, plants possessing the succulent syndrome are thought to have evolved to adapt to arid climates, because they possess modifications that increase their water use efficiency. Here we study the evolution and the ecological nature of the succulent CAM syndrome at a global scale by analyzing the climatic niches of succulents within the Caryophyllales, testing the hypothesis of a climatic niche specialization by comparing them with their non-succulent, non-arid adapted relatives. We assembled and carefully curated a worldwide dataset of 5447 species in 28 families, and analyzed the current and evolutionary trajectories of climatic niches with an array of statistical methods including ecological niche modeling, phylogenetic regression and divergence dates estimation. Our results confirm the Core Caryophyllales tend to inhabit drylands probably since their origin in the Early Cretaceous. However, the succulent syndrome appeared later with some lineages diversifying profusely afterwards. The climatic niche of succulents is not differentiated from their non-succulent relatives, but narrower, and contained within the non-succulents’, showing no relationship with extreme conditions such as high aridity or temperatures. Our results support alternative interpretations of the origin of the CAM syndrome and the ecological significance of succulence, as well as the prolific radiation of richest lineages.

**Highlights:** The climatic niche occupied by succulent CAM plants is not different from their non-succulent relatives. Estimated dates and character reconstruction suggest CO2 scarcity as the evolutionary pressure under these plants originated.

## Introduction

Succulent plants evolved the most dramatic modifications observed in the plant kingdom, which occurred at all organismal levels (Gibson and Nobel, 1986; Lüttge, 1987; Gibson, 1996; Mauseth, 2006). They achieved a complete morphological and anatomical transformation of their tissues and organs to store water, developed a novel metabolic pathway, modified developmental programs (e.g. loss or reduction of organs), or changed the physiological regulation of certain processes (e.g. patterns of stomata opening). All modifications present in succulents are highly interrelated, and together they are known as the “succulent syndrome” (Ogburn and Edwards, 2010; Griffiths and Males, 2017). Although different succulent lineages show variations in the succulent syndrome, several traits are common among all succulents, like the reduction of certain organs together with an increase in the size of others that store water, the development of large vacuoles, the presence of thick cuticles, low stomatal densities, etc. (Gibson and Nobel, 1986; Mauseth, 2004 *a*,*b*; Ogburn and Edwards, 2010; Griffiths and Males, 2017; Fradera-Soler et al., 2022).

One keystone element in the succulent syndrome is the Crassulacean Acid metabolism (CAM), a type of photosynthesis that is highly efficient in the use of water. Important efforts are being made to understand and manipulate the CAM succulent strategy in order to develop resilient crop systems to future climate change scenarios of increased heat and drought (Borland *et al*., 2009; Borland *et al*., 2015; Yang *et al*., 2015; Mason *et al*., 2015; Leverett et al. 2021; Moreno-Villena et al., 2022). Systems biology approaches and physiological models have been proposed (DePaoli *et al*., 2014; Yang *et al*., 2020), and genomic sequences have been generated for several species to decipher the CAM genetic blueprint (Hartwell *et al*., 2016). Given their high water use efficiency (WUE), succulent-CAM plants have been typically considered as iconic examples of organisms well adapted to dry and hot environments (Gibson, 1996; Herrera, 2009; Kluge and Ting, 2012), and are recurrently used as examples of plant adaptations in evolution or plant biology text books (Niklas, 1997; Lack and Evans, 2001 pp. 122; Futuyma, 2005 pp. 118; Stern *et al*., 2006 pp. 276; see references in Evans *et al*., 2014; Fradera-Soler *et al*., 2021), as well as in more specialized literature (Edwards and Donoghue, 2006; Nyffeler *et al*., 2008; Ogburn and Edwards, 2010; Holtum *et al*., 2016), in particular in relation to the CAM metabolism (Ting, 1989; Keeley and Rundel, 2003; Edwards and Ogburn, 2012). However, the ecological complexity of the succulent strategy is often overlooked, as well as the diversity of environments these plants inhabit. The dryness of arid regions has led to the false assumption that desert plants possess a physiological resistance to drought, however, ecophysiological investigations show the view is incorrect and the water supply of desert plants is not so poor (Walter, 1973). This is particularly for succulent CAM plants, since this strategy is not necessarily more abundant in the extreme desert environments (Griffith and Males 2017).

The succulent CAM strategy evolved several independent times along the evolution of angiosperms, with succulent species present in about 400 genera distributed in around 36 families (Silvera *et al*., 2010). In general, the early-diverging members of succulent lineages have less degrees of succulence or are not succulent at all, sometimes still possessing a C3 type of photosynthesis or facultative CAM (Rayder and Ting, 1981; Applequist and Wallace, 2000; Klak *et al*., 2003; Edwards and Donoghue, 2006; Edwards and Diaz, 2006; Nyffeler and Eggli, 2010a; Hernández-Hernández *et al*., 2011). Additionally, these early divergent members tend to have broader distributions, while succulent species within derived clades tend to concentrate in localized regions, indicating a possible evolutionary trend towards a niche specialization that occurred relatively recently (Axelrod, 1972). Moreover, it has been suggested that lineages showing the succulent syndrome synchronously evolved convergent solutions to a similar problem of water scarcity in different regions of the world, indicating a parallel adaptive response to a global climatic trend (Arakaki *et al*., 2011; Edwards and Ogburn, 2012). However, the ecological and evolutionary context of the origin and evolution of the succulent CAM syndrome is still insufficiently understood, and detailed data about the evolution of succulence in relation to arid climates (like high temperature and low precipitation) are needed to better understand and manipulate this plant strategy.

The angiosperm order Caryophyllales, including ca. 38 families, is an excellent system to study the evolution of arid-adapted succulent lineages and their strategies in an evolutionary and ecological context. Within the order, a number of families evolved extreme succulence, and some of these lineages show the highest diversification rates observed in the plant kingdom, like Cactaceae or Aizoaceae (Klak *et al*., 2004; Magallón and Castillo, 2009; Hernández-Hernández, et al. 2014). Here we analyze the ecology and evolution of succulent Caryophyllales in terms of their habitat and climatic niche. With an assembled database of climatic data extracted for 201,734 revised locality records corresponding to 5447 species within 28 families included in the Core Caryophyllales, we investigate the relationship between the presence of succulence and environmental conditions. We focus our analyses on the climatic components of aridity: low precipitation regimes, high precipitation seasonality and high temperatures; plus aridity indexes, potential evapotranspiration and natural fire regimes. By analyzing this global dataset, we evaluate if succulent plants occupy a distinct climatic niche space. Particularly we test whether they show a climatic specialization towards drier and warmer environments in relation to their non-succulent, temperate or tropical relatives. We analyze the evolutionary trajectories of climatic niches within the group using phylogenetic comparative methods, and evaluate if there is an evolutionary tendency towards inhabiting arid and extremely arid conditions in derived succulent clades. Finally, we test for the convergence of independent succulent lineages towards the occupation of similar climatic niches in different regions of the world using ecological niche models (ENMs).

Our statistical analyses show succulents’ climatic niche is narrower than their non-succulent relatives, it is contained within the non-succulents one, and does not show a statistically significant differentiation towards drier and warmer regions of the Core Caryophyllales global climatic niche. Moreover, some representatives of the non-succulents occur in the ranges of the niche space with the highest temperatures and lowest precipitation. ENMs were able to predict the geographic patterns of succulents’ diversity. However, the richest succulent lineages that are distributed in different regions of the world do not inter-predict each other, suggesting that they occupy different climatic niches. Our results show that succulents are not more abundant towards the extreme arid space of the climatic niche, which might suggest a possible trade-off in these lineages of climatic niche versatility for water use efficiency, perhaps facilitating versatility in reproductive strategies (Herrera, 2009). Studying the ecology of succulent CAM plants in an evolutionary context provides important information to better understand the origins and the ecophysiological implications of their complex WUE strategies.

## Materials and Methods

### Phylogenetic analyses

We followed two different approaches for the phylogenetic analyses. First, we assembled a combined matrix of DNA sequences with the help of the PhyLoTA browser (Sanderson *et al*., 2008) for representative taxa of the 38 families included in the order Caryophyllales (*sensu* Stevens, 2016), plus 9 species to represent outgroup families (Cuénoud et al., 2002; Stevens, 2016). We selected the *matK*, *atpB*, *rbcL* and the 18S genetic regions given their good representation in the study group. We focused our sampling in the Core Caryophyllales, and included all genera and species with DNA sequences available for at least two of the four genetic regions considered. The matrix includes 150 ingroup and 11 outgroup taxa (161 data matrix, Appendix S1, Supplementary Information). The length of the aligned matrix is 7336 bp (Appendix S2). Following a second approach, we obtained another phylogeny by using a pruned database obtained from the original data in Smith *et al.,* 2018 to achieve 721 taxa (721 taxa dataset, Appendix S2). This matrix is 12,714 bp and includes the plastid markers *matK*, *ndhF*, *rbcL*, *trnH-psbA* spacer, and *trnL-trnF* spacer; as well as the nuclear ribosomal internal transcribed spacers (ITS) and the *phyC* gene (Smith *et al*., 2018). In both cases, we performed phylogenetic analyses under default options using RAxML, including 100 ML bootstrap analyses to evaluate node support. The estimation of divergence dates was performed using in BEAST v2.4.7. Information on analyses details and calibrations are available in the Extended Methods section of the Supplementary Information.

### Locality data, habitat description, and climatic variables

Occurrence locality data for all available species within the families included in the Core Caryophyllales (see Table S1) were downloaded from the Global Biodiversity Information Facility (GBIF) database (https://www.gbif.org) using family key word searches from September 2017 through January 2018. We followed the currently accepted classification and families available in the Angiosperm Phylogeny Website or APW (Stevens, 2017). Data found in GBIF is partially curated and prune to identification and geo-localization errors. Given the size of our database (5447 records) we were not able to corroborate species identification and distribution for each record, and instead we performed a semi-automatic curation process. For this reason, we acknowledge that there might be sources of error inherent to the nature of our database, but we hope that the size of the data analyzed and scope of our questions provide a reliable signal for succulent plants as a group. Our curation strategy was implemented mostly at the genus level. In summary, we revised the taxonomical names, removed specimens not determined at the genus level, removed *ex situ* specimens that were cultivated or growing in gardens and living collections, and confirmed distribution by comparing with data from APW (Stevens, 2017) and Kubitzki (1993). The strategy is fully explained in the Extended Methods. Since it would have been extremely time-consuming to obtain and analyze information for each species in the database (5447 spp), and because for most species the information is not available or non-existent, we classified species following the information available for the genus they belong according to their habitat and degree of succulence. We classified species within genera according to the biomes they inhabit as *arid*, *temperate*, or *widespread* (see Extended Methods). We also classified the habitat of each locality according to Noy-Meir (1973) classification of desert ecosystems based on annual mean precipitation values: (1) extremely arid (less than 100mm), (2) arid (from 100 to 250mm), (3) semiarid (from 250 to 500mm) and non-arid if more than 500mm. In addition, we classified habitat following CGIAR-CSI aridity index values (AI, https://cgiarcsi.community) as following: hyper arid when AI<0.03, arid when 0.03<AI<0.2 semi-arid when 0.2<AI<0.5, dry sub-humid when 0.5<AI<0.65 and humid when AI>0.65. Finally, we classified species in our database according to the succulence degree of the genera where they belong as *succulent*, *fleshy* and *non-succulent*. Details for classification criteria can be found in the Extended Methods and Appendix S1.4.

Environmental data for each locality was obtained from the CHELSA climatic database (Karger *et al*., 2017), which contains climatic data layers at a spatial resolution of 30 arc seconds (∼1 km resolution). We only used the variables hypothesized as the more important controlling factors for organisms in desert ecosystems, i.e. those reflecting extreme temperature ranges, low precipitation and seasonality (Noy Meyr, 1973; McGinnies, 1979): Annual mean temperature (Bio1), maximum temperature of warmest month (Bio 5), minimum temperature of coldest month (Bio6), annual precipitation (Bio12), precipitation seasonality (Bio15), precipitation of wettest quarter (Bio16), and precipitation of driest quarter (Bio17). Additionally, for each locality we obtained data on potential evapo-transpiration (PET), aridity index (AI) (Zomer *et al*., 2008), soil-water balance (alpha index; Trabucco and Zomer, 2010), and fire regimes from the Global Fire Atlas (Giglio *et al*., 2018; Andela *et al*., 2019). To evaluate the possible effect of other variables not selected, we repeated our analyses with all 19 variables including in the CHELSA climatic dataset.

### Evolutionary and statistical analyses

The Caryophyllales phylogenies were pruned to obtain a tree including one representative taxa per family within the Core Caryophyllales. *Rhabdodendron* (Rhabdodendraceae) was kept as outgroup (Appendix S2). The percentage of succulent species was estimated for each family in the database, as well as the median values for each climatic variable (Appendix S1.2). We evaluated a possible relationship between succulence or degree of succulence (dependent variable) and the variables determining arid climatic niches (independent variables, i.e. aridity index, habitat, and bioclimatic variables) taking into account for phylogenetic relationships using PGLS regression (Martins and Hansen, 1997), performed using the R package *caper* v0.5.2 (Orme, 2013) following standard practice. We also performed PGLS analyses to test for a relationship amongst the presence of succulence or the degree of succulence within families and their diversification rates. Net diversification rates for each family were estimated using the standard method-of-moments estimator for stem-group ages (Magallón and Sanderson, 2001), using the R package *geiger* v2.0.6 (Pennell *et al*., 2014). We analyzed the evolutionary trajectories followed by the studied variables amongst families by constructing traitgrams with the *phenogram* function in the R package phytools (Revell, 2012). We used the *fastAnc* function to find the maximun-likelihood estimate of ancestral character states over the phylogeny for each variable.

Finally, we explored the behavior of climatic variables regarding locality and groups of species using different statistical approaches. We first compiled an aggregated database by summarizing climatic variables by median values per species. Weighted principal component analysis (WPCA) was used to reduce dimensionality on this aggregated database. Given the large number of species, we used density estimators instead of ordinary scatter plots, which generally become obscured because of overplotting. We also examined the potential of climatic variables to predict categorization into accepted degrees of succulence via logistic regressions, and summarized the potentiality of such classification in ROC (Receiver Operating Characteristic) curves (see details in the Extended Methods).

### Succulent potential richness areas and climatic convergence

By adding more data and performing a more detailed curation, we assembled another improved database for lineages showing the highest succulence within the Caryophyllales. This database allowed us to use ecological niche modeling (ENM) to (1) calculate the geographic areas with the highest potential of occurrence of succulent species for the lineages selected, and (2) to evaluate the possible convergence among succulent lineages that inhabit different regions of the world, towards similar climatic regimes. In other words, we aim to test whether hotspots of succulents’ diversity in different geographic regions show similar climatic conditions. For this we selected the following lineages to whom size or geographic distribution allowed to achieve a better curation of the georreference data: Cacteae and Pachycereeae tribes within Cactaceae (the globose and columnar cacti of North America), the Didiereaceae family, the Ruschioideae subfamily in Aizoaceae as well as Anacampserotaceae and the genus *Portulacaria* (Portulacarioideae, see Bruyns *et al*., 2014, Figure 5). Although there are many succulent species of Cactaceae distributed in South America, due to the scarcity of reliable locality data and geographic range reports, we did not include them in our study. Locality data for members of the selected groups was obtained from different sources, and the natural distribution was revised more carefully (at the species level) by using specialized literature (see details about occurrence records and curation strategy in the Extended Methods).

We estimated ENMs for every species on these datasets by using MaxEnt (Philips *et al*., 2006), with the environmental variables described in the previous section but with a downscaled resolution of 2.5 arc minutes (∼ 5 km at the equator). These predictions were stacked by summing the individual continuous predictions at the local scale to estimate the areas with the highest potential richness of succulent species (potential richness areas, PRA). We evaluated the climatic convergence among PRAs of the different lineages by projecting and stacking the individual ENMs of species within each lineage at the global scale, to generate projected potential richness areas (PPRAs). When doing these projections, we restricted them to extrapolate the niche models within the range of training values. In case of convergence, PRAs and PPRAs of different lineages should be similar or at least would inter-predict the distribution of the succulent lineages amongst different regions of the world. Convergence was evaluated visually, and analyses were performed in the statistical software R using the packages *raster* (Hijmans, 2015) and *letsR* (Vilela and Villalobos, 2015).

## Results

### Climatic distribution of succulent and non-succulent Core Caryophyllales

Our assembled localities database includes 28 families, with 574 genera and ca. 5447 species, comprising 201,736 entries with georeferenced locality data (Figure 1A). These taxa include 89% of genera and 66% of species within the Core Caryophyllales, which inhabit a wide range of habitats and ecosystems (Figure 1A). Both succulent and non-succulent Caryophyllales are broadly distributed across all continents, except for Asia where they are scarce. However, the maps indicate that the highest diversity of succulent and fleshy records is concentrated in South Africa, arid regions of North America, Brazil, and Australia (Figure 1B-C). This distribution pattern is also congruent with hotspots of distribution of succulents in other lineages (e.g., *Agave*, *Aloe*, *Euphorbia*, etc.). Although it can be seen that the Core Caryophyllales tend to distribute in arid or semiarid regions across the world, they are very scarce in the arid regions of Northern Africa (Saharan Desert and surrounding areas), the Arabian and Syrian Deserts, or in arid regions of Asia (Gobi Desert), where the extreme arid regions of the world are found.

**Figure 1.**
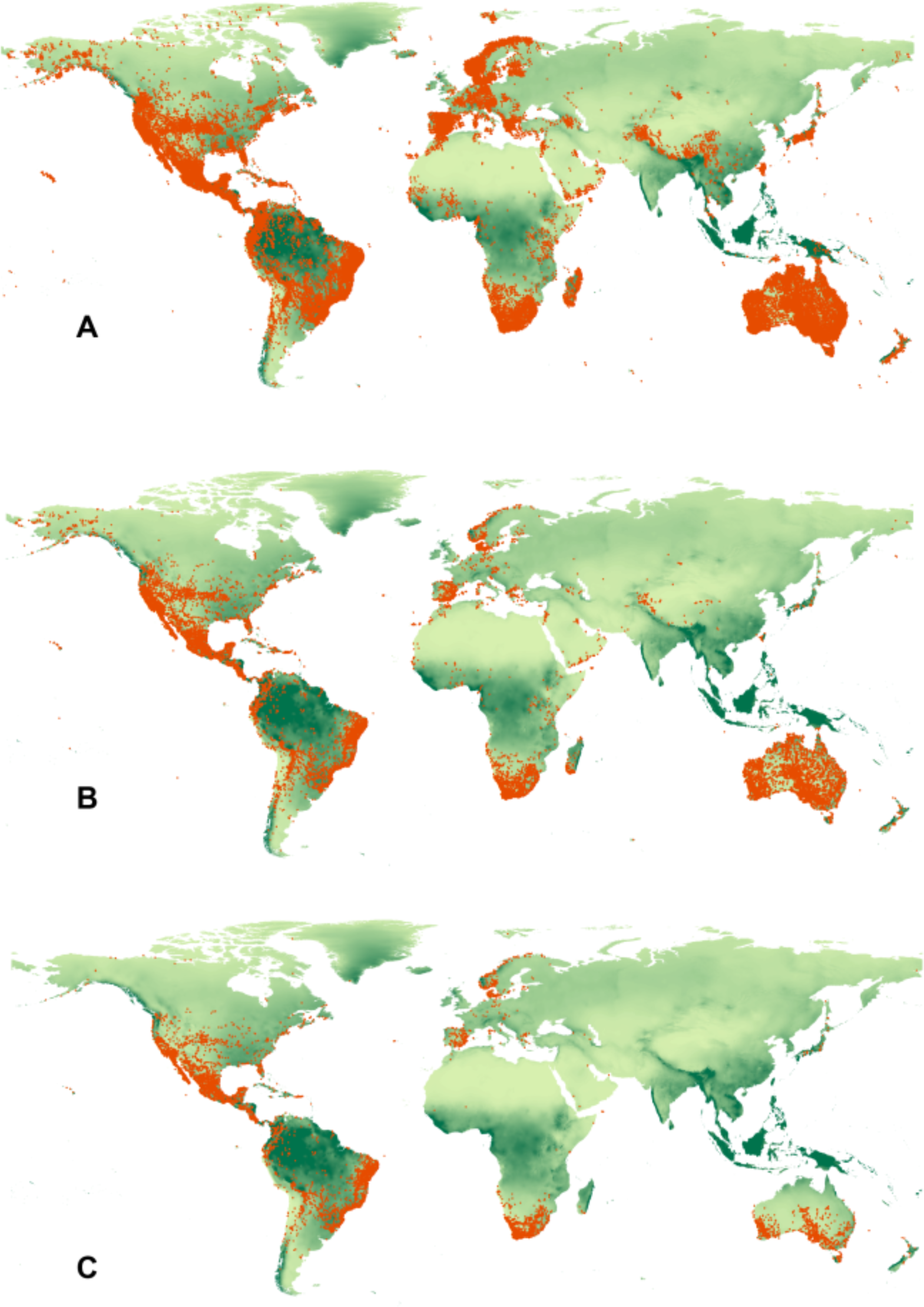
Geographic distribution of locality records in our database (A) for succulent and fleshy species (B), and only for succulent (C).

Annual mean precipitation ranges amongst all species are large, the extreme values being 0 (e.g. *Acyranthes*, *Althernantera* records for Amaranthaceae in extremely arid regions in Libya) and >7600 mm (e.g. *Pereskia*, *Amaranthus* and *Phytolacca* records in tropical humid Colombia). According to our estimates, most species are widespread (inhabiting more than one type of habitat or biome), while almost 40% inhabit arid, 9% inhabit tropical and 5% are located in temperate biomes (Table S3). Most families include species with records present in all the different categories of aridity as well as non-arid environments (Table S1-S2). The exception is Barbeuiaceae, which is a monotypic family inhabiting exclusively humid environment. Following Noy-Meyr’s classification, most of the records in the database correspond to non-arid (55.85%), followed by semiarid (28.7%), arid (13.4%) and 2.05% are in extremely arid regions. Following the CGIAR-CSI AI (Aridity Index) criteria, 63.74% of all our registered localities fall within dryland regions, while 36.26% fall within humid regions (Figure 2). From all our records, 36.22% correspond to fleshy or succulent plants. These belong to 53.9% of species in our database. In addition, 15.43% of records, belonging to 36.3% of species in the database are highly succulent species (not fleshy). The distribution of succulent and fleshy plant records in our database per AI category is very similar to the distribution of all records in the entire database (including non-succulents, see Figure 2). For most climatic variables analyzed, non-succulents, succulents and fleshy plants (low succulence) show similar median values (Figure 3).

**Figure 2.**
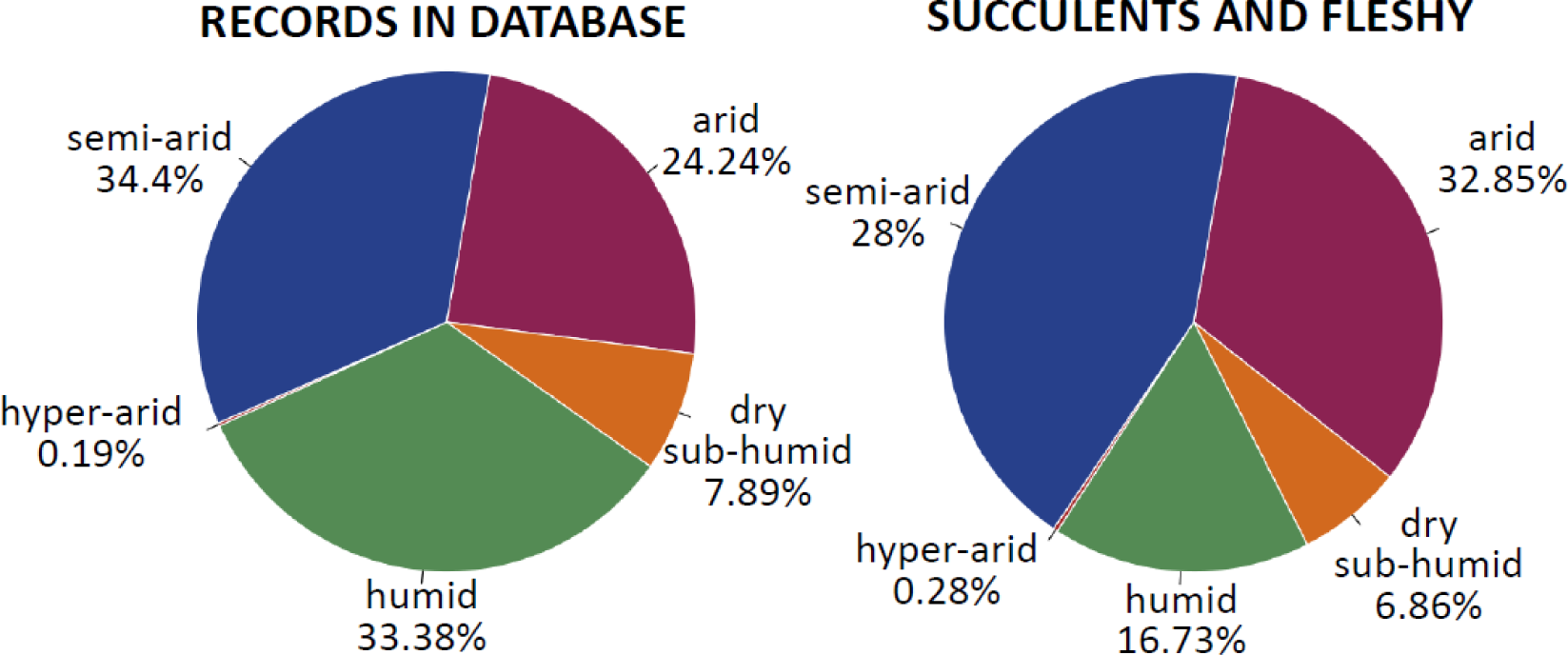
Distribution of all records in database (left) and only succulent and fleshy records (right) per aridity index categories (CGIAR-CSI criteria).

**Figure 3.**
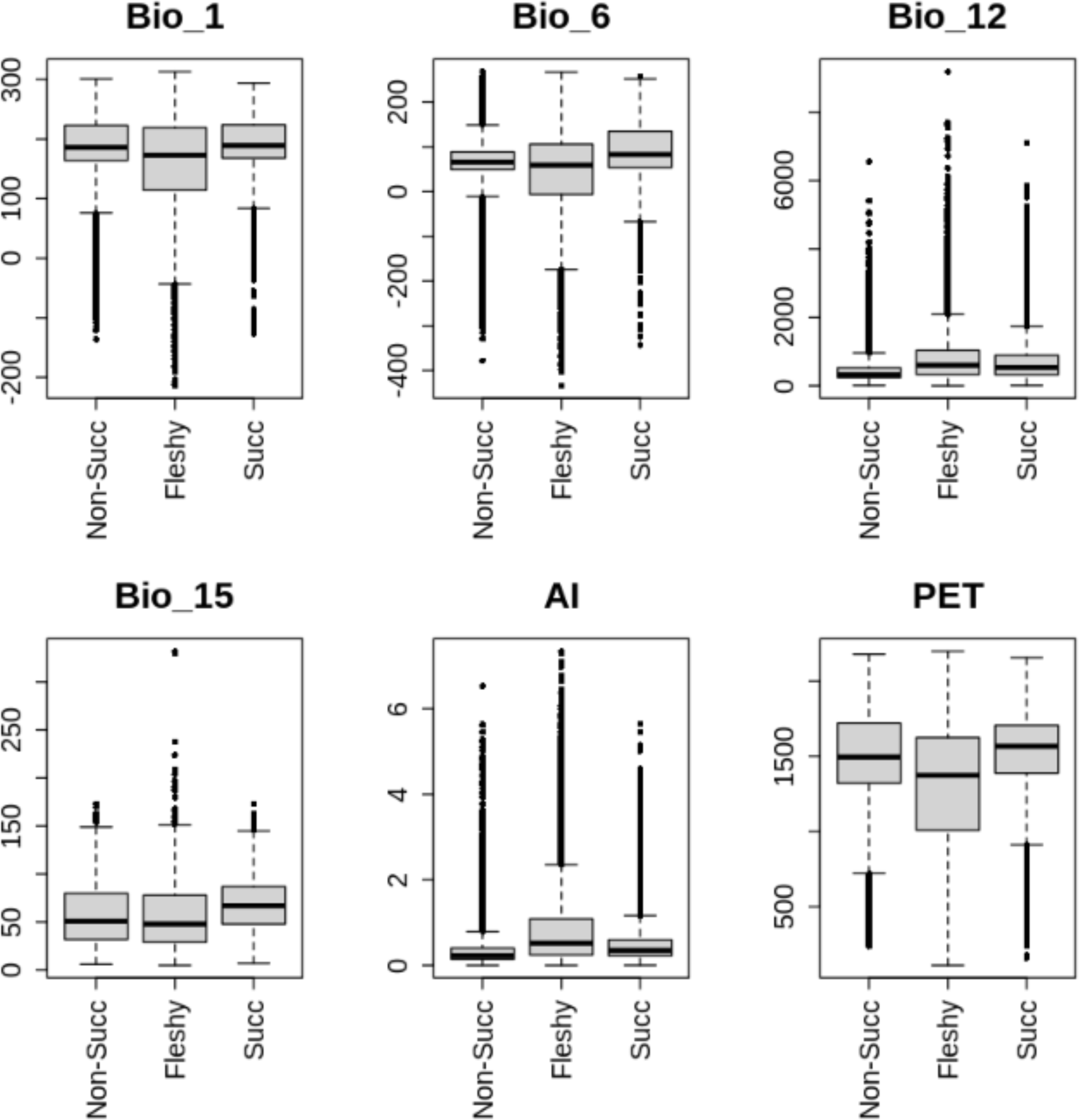
Empirical distributions for some of the environmental variables considered in this study for species classified as non-succulents, fleshy and succulent within the Core Caryophyllales: Bio1 (annual mean temperature), Bio6 (minimum temperature of the coldest month), Bio12 (annual precipitation), Bio15 (precipitation seasonality), AI (aridity index), and PET (mean potential evapo-transpiration per year).

Before conducting WPCA analyses we examined the potential of variables to predict the degree of succulence by means of logistic regression using ROC curves (Figure S4). We found that AI, alpha (soil water-balance), and fire regime variables do not contribute significantly in predicting succulence when added to other variables, and we excluded them from further analyses. AI and alpha are likely redundant given that bioclimatic variables and PET are involved in their estimation (https://cgiarcsi.community). To account for a differential representation of records for species within our database we performed WPCA at the species level. The first two components, WPC1 and WPC2 explain 47.89% and 34.21% of the variance, and have a clear interpretation (Figure 4a). In general, they correspond to temperature and precipitation variables respectively, however the temperature component (WPC1) also includes PET and precipitation seasonality (bioclimatic variable 15, see Figure 4a). Figure 4b shows WPC1 and WPC2 displayed on a scatter plot. As it is difficult to discern structure due to severe overplotting (Figure 4b), we used density functions, elaborating contours that enclose highest density regions at two levels: 75% (Figure 4c-d) and 95% (Figure 4e-f). The visual examination of figures show that climatic niches of succulents and non-succulents are not disjoint in the climatic space, since the succulents and fleshy niches are narrowly concentrated within the Core Caryophyllales (Figures 4c-f). Since the classification of taxa as fleshy might be subjective, we repeated all analyses considering them either as succulent or non-succulent (see Figure S5). Results are similar to the explained above. To evaluate if we failed to include a climatic variable that might be relevant for the hypotheses tested, we repeated the WPCA analyses including all 19 CHELSA variables, and visually compare our graphs with the WPCA built with only selected variables. Results with all CHELSA variables are highly similar, indicating the relevance of the variables selected (Figure S1, Supplementary Material). We further discuss our results following the selected variables described in the Methods section.

**Figure 4.**
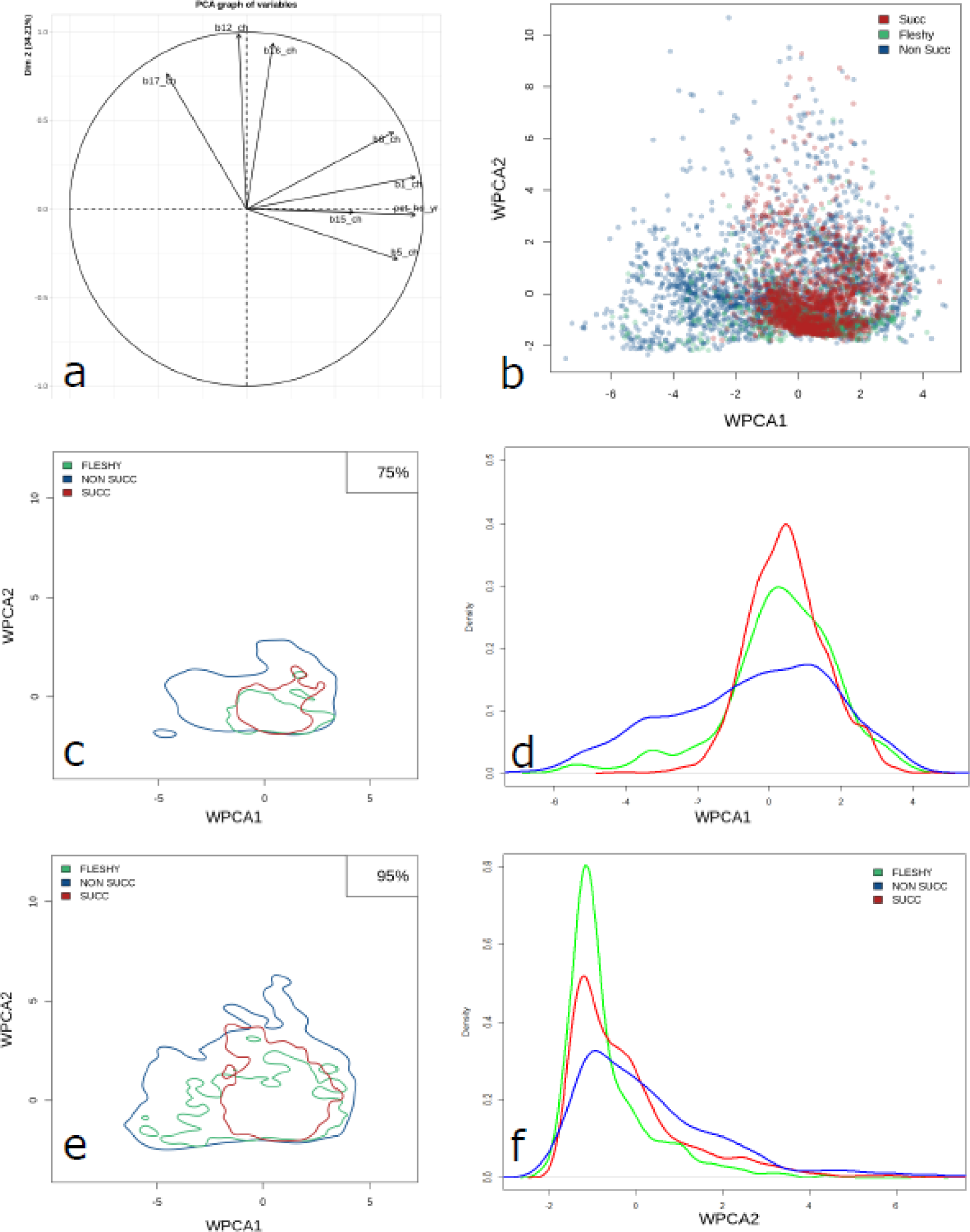
Results of the WPCA analyses. (a) biplot of weights of the first two WPCA’s, (together account for 82.77% of variability), (b) plain scatter plot of WPCA1 and WPCA2, where each point represents a single species, (c) contours of 75% highest density regions (HDR) per plant group, (e) contours of 95% HDR’s, (d) marginal densities of WPCA1 by group, and (f) marginal densities of WPCA2 by group. Blue: non-succulent; green: fleshy; red: succulent. Contours and marginal densities show the values where the majority of observations are located in the WPCA.

### Phylogenetic analyses, estimated dates and evolution of climatic niches

The topology and phylogenetic relationships we obtained for the Order Caryophyllales (Appendix S2, Figures S2 and S3 Supplementary Material), both with the 721 and 161 taxa data sets are highly similar amongst them and to the most recent phylogenies reported for the group with a more extensive taxonomic sampling and large amounts of DNA sequence data (Walker *et al*., 2018; Yang *et al*., 2018). In general, relationships are well supported by bootstrap values (Appendix S2, Figures S2 and S3). The Core Caryophyllales is a strongly supported monophyletic clade (phylogenetic trees in Appendix S2). At the family level, the taxonomic groups with the highest diversification rates are Cactaceae and Nyctaginaceae (0.15 and 0.1 sp/Myr) respectively. Their diversification rates are even higher than other families within the Order which include a higher number of species (for example, Amaranthaceae or Caryophyllaceae, see Appendix S4).

Our dating analyses reveal similar dates to previous reports (Appendix S1.3). However, our dating analyses with 161 taxa matrix and results in Arakaki *et al*. (2011) are younger.

Results with the 721 taxa matrix, dates obtained by Ramírez-Barahona *et al*., (2020), and the PL analyses results of Smith *et al*., 2018 are similar, but confidence intervals are wide in most cases in the first two BEAST analyses (Appendix S1.3). Our results indicate the Core Caryophyllales originated around 127/128 Mya during the Early Cretaceous (mean heights obtained in both the 161/721 taxa matrix analyses). However, our results indicate families diversified much later (mean stem age for Core Caryophyllales families is 55.7 Mya), and started a proliferation more recently, around 25.7 Mya (Appendix S1.3). The estimated dates suggest the groups with the highest proportion of succulent species (i.e., Aizoaceae, Didiereaceae, Cactaceae, Anacampserotaceae, Halophytaceae, Kewaceae and the Portulacarioideae) originated each at different, non-overlapping time frames.

However large confidence intervals as well as the variation found among estimates of different analyses make it difficult to conclude a precise and accurate time frame for the evolution and diversification of clades. Our estimates suggest Kewaceae originated first, around 58.5 Mya. However estimated dates range from 44.8 to 99.7 Mya. Aizoaceae might have originated around 49.6/69.1 Mya, but started diversifying ca. 25 My later.

Portulacarioideae (*Portulacaria* + former *Ceraria*) originated later, perhaps around 35.6/43 Mya, but we estimated a very recent diversification much later, perhaps at 1.7/9.4 Mya. Although Cactaceae, Didiereaceae and Anacampserotaceae originated at similar time frames (stem dates around 25.2/44.18, 39.7/42.9, 25.2/44.18 Mya according to median estimates from the 161/721 taxa matrix analyses), crown estimated dates suggest they started diversifying at different times, particularly the highly succulent clades within them (see Appendix S1.3).

PGLS analyses at the family level show a significant relationship of fleshy and succulent prevalence in the family with semi-arid conditions as well as two bioclimatic variables related to precipitation (bio 12 or annual precipitation and bio 16 or precipitation of the wettest quarter) however with low r-squared values. There is not a significant relationship with variables related to elevated temperatures (such as the maximum temperature of the warmest quarter, or bio 5) or extremely dry conditions (such as arid or hyper-arid classifications); potential evapo-transpiration or precipitation seasonality. There is only a slightly significant relationship with annual mean temperature (bio 1), or precipitation of the driest quarter (Appendix S1.5). Interestingly, the presence of fleshiness or succulence is not correlated with diversification rates, possibly because some families with very low number of species (Kewaceae or Halophytaceae) were classified as having 100% of their members as succulent.

Traitgrams and ancestral climatic conditions were obtained for bioclimatic variables 1, 6, 12, 15, AI, and PET (Figure S6 and Appendix S1.6). Results show mild temperatures for the ancestor of the core Caryophyllales (ca. 20°C annual mean temperature, 11° minimum temperature of coldest month), annual precipitation corresponding to non-arid environments (923mm, Noy-Meyr classification), and aridity index corresponding to dry sub-humid regions (AI=0.57, CGIAR-CSI classification). When mapping the ancestral estimated conditions into modern world climates (Figure S7) we see that although there is some coincidence with semiarid lands, they are more coincident with dry sub-humid conditions.

The projections of the ENMs showed accurate predictions of the known distributions of the species with 356 out of the 430 species showing reliable model performance (average TSS values above 0.75). After stacking the individual species predictions to obtain the PRAs per succulent lineage, we observed ENMs were able to predict the known geographic hotspots of succulents’ diversity in the world (Figure 5). However, ENMs of succulent lineages were not able to geographically inter-predict, suggesting different climate regimes per PRA. Members of the Anacampserotaceae, that have the largest distribution amongst the six lineages, partially overlap with Cactaceae in North America and with Ruschiodeae in South Africa, and it is the only example with inter-prediction between lineages.

**Figure 5.**
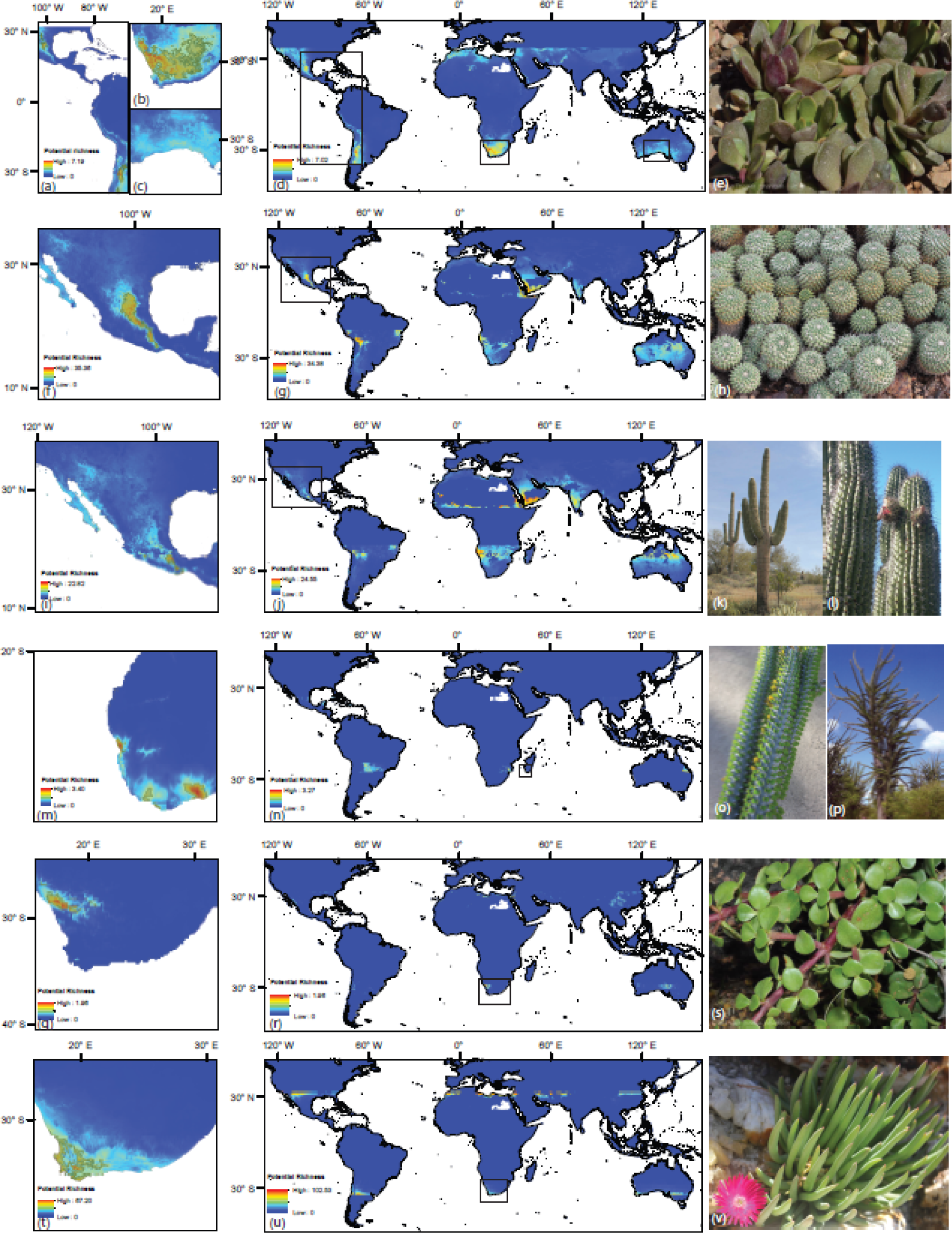
Ecological niche model predictions of the six Caryophyllales lineages showing high levels of succulence: (a - e) Anacapserotaceae, (f - h) Cacteae, (i-l) Didiereaceae, (m - p) Pachycereae, (q - s) Portulacarioideae and (t - v) Ruschiodeae. Maps of potential richness areas per lineage in their native range are shown in the left panel, whereas the projection of their climatic conditions into the world space are shown in the central panel. The species depicted in the right panel are: (e) *Anacampseros rufescens* *, (h) *Mammillaria compressa* ^§^, (k) *Allaudia procera* ^§^, (l) *Allaudia procera* *, (o) *Carnegiea gigantea* ^§^, (p) *Stenocereus thurberi* ^¶^, (s) *Portulacaria afra* *, (v) *Cephalophyllum regale* ^§^. Photograph credits: (*) © 2014 Jan Thomas Johansson, (^§^) I. Loera, (^¶^) T. Hernández-Hernández.

## Discussion

### Climatic niches of succulents are not very different from their non-succulent relatives

It has been estimated that dry climates are the most extensively distributed over the earth, covering around 26% of the continental area (McGinnies, 1979). There’s a prevailing idea that the origin of these water-stressed environments conferred the adaptive pressure under which several plant lineages rapidly evolved a variety of strategies and specialized structures that enabled them to withstand periods of severe drought (Stebbins, 1952; Axelrod, 1972; Gibson, 1996; Ward, 2016). Among those strategies, succulence has been regarded as the one suggesting more evidently a correlation with climatic variables such as precipitation and temperature (Von Willert *et al*., 1992; Gibson, 1996; Ward, 2009). However, a detailed characterization of the climatic niche of succulents has never been provided, and the relationship among the morphological prevalence of succulence and climatic variables has been tested only in few of studies. Evans *et al*., 2014 showed that in the *Euphorbia* of Madagascar, water storage is associated with the two components of aridity: temperature, and precipitation, but in the South African *Crassula* genus, microhabitat preferences influence more the morphological evolution towards leaf succulence and xerophytic characters than aridity at a macroclimatic level (Fradera-Soler *et al*., 2021). The relationship amongst the prevalence of CAM and climatic conditions has been addressed more often, however either in non-succulent taxa or without considering succulence as a primary component of the CAM syndrome. Studies in the non-succulent Eulophiinae orchids (Bone *et al*., 2015), non-succulent Bromeliaceae (Males and Griffiths 2017b; Males, 2018 *a*, *b*), or the Agavoideae (Borland *et al*., 2015) suggest a concordance with the occurrence of CAM and arid conditions as well as with functional types associated with climatic types.

A preliminary analysis of climatic data for 883 species within the Portullugo clade (Caryophyllales), showed that taxa with C4 and CAM metabolism occupy a similar climatic space, suggesting already that both pathways evolved in this group under the same set of environmental conditions (Edwards and Ogburn, 2012; Edwards et al., 2019). Unfortunately, an unambiguous determination of a species as a CAM or C4 requires experimental approaches and anatomical observations, often under controlled conditions, that are currently impossible to achieve for a large number of species. For that reason, we focused our study on succulence, as an adequate proxy to CAM. Although the CAM has been found in non-succulent plants, succulence has been shown as a requisite for obligate CAM (Niechayev et al., 2019; Griffiths and Males, 2017), and to our knowledge, there have not been succulents reported as being non-CAM.

In this study, we explicitly test the relationship of succulence and climatic variables at a global scale, using a taxonomic group large enough to perform robust statistical analyses and allowing a comparison with evolutionary related non-succulent, non-arid adapted groups. The complexity of the succulent niche in relation to water availability has already been pointed out. Early analyses of botanical data showed that although succulents show excellent WUE, they dominate places where precipitation is small but offered regularly (Ellenberg, 1981). Under irregular conditions, several species of non-succulent shrubs or trees are superior, and especially tall stem succulents are mainly favored by low but regular annual precipitation (Ellenberg, 1981). Our results confirm these early observations, and provide a modern perspective with improved datasets that allows us to better understand the origin, evolution and ecological significance of succulence, and associated CAM strategy.

The distribution of succulent species over different environments is not significantly different from the data for the entire Core Caryophyllales, suggesting similar niches both for succulents and non-succulents (see Figure 2, Appendix S1). Although we estimate almost half of the species within the Core Caryophyllales are succulent or fleshy (53.9%, Figure 2, Table S1), 23% of these succulents inhabit humid environments and less that 1% survive the hyper-arid regions (Figure 2), showing the versatility of the succulent strategy, and that the idea that succulence is an adaptation to aridity or extreme aridity might be a reviewed. PGLS analyses results do not show a strong correlation among prevalence of succulence and variables indicating extreme aridity, but although with low significance and r-squared values, they do show a relationship with semi-arid or dry-subhumid conditions, that might be the result of a general tendency in all members of the group (see PGLS results in Appendix S1.5). The WPCA also shows that the climatic niche distribution of succulents overlaps with the non-succulents, and that succulents do not show a tendency towards the extreme section of the climatic niche with low precipitation, low humidity and higher temperatures. In fact, whereas fleshy and non-succulent species are present in all regions of the climatic niche space, it is the non-succulents that are able to reach the extreme conditions (both towards the humid and towards the hot and dry, see Figure 4).

Although we didn’t find a relationship among succulence and diversification rates, important succulent lineages have shown to have some of the highest diversification rates amongst vascular plants (e.g. *Agave*, Cactaceae clades or the Ruschioideae; see below). What could be the ecological meaning of the succulent strategy in addition to overcoming environmental water deficits? By observing the sharp concentration of succulents in the niche space (red and green lines in Figure 4d and f) we see that succulents might not be as prevalent towards the extreme conditions of the niche space they occupy (both towards de humid or towards the dry regions). A major difference of the succulents’ life strategy is their carbon and water balances (Von Willert *et al*., 1990). In succulents, the possession of water reservoirs allows them to afford energy investments in growth and reproduction even during the dry season. It is known that several succulents, both in the old and the new world, flower during the dry season (Von Willert *et al*., 1990). As it has been suggested before (Herrera,2009), the development of a water storage strategy, in addition to provide advantages to survive water scarcity, can help increase the reproductive versatility of succulents, allowing these lineages to explore the reproductive niche space (e.g. phenology or pollination syndromes) in time and space more broadly than non-succulents. In this scenario, succulence and the operation of CAM may be associated with an increase in fitness by increasing reproduction under drought (Winter and Ziegler, 1992; Herrera, 2009).

### The Core Caryophyllales show an ancient trend towards arid environments

Although succulents are not more abundant in extremely or hyper-arid regions, a high proportion of records and species within the Core Caryophyllales (both succulent and non-succulent) are present in drylands in general (66.62%, Figure 2), indicating a clear trend to inhabit water stressed environments in the group. Interestingly, the earliest diverging member within the Core Caryophyllales, the monotypic family Simondsiaceae (*Simmondsia chinensis* or jojoba), an evergreen non-succulent shrub, is the lineage showing the highest abundance in hyper-arid regions, inhabiting the most extreme conditions. This species is also present in all remaining climatic regimes, showing the versatility of this ancient and depauperated lineage (ca 128 Mya Appendix S1.3).

We estimated a very ancient origin for the core Caryophyllales (ca. 128 Mya, chronograms in Appendix S2) during the Early Cretaceous. There are sedimentary evidences for global increases in aridity from the Cretaceous (Ziegler *et al*., 2003). There is also evidence of aridity in this period coming from macrofossils exhibiting extreme xeromorphism and clear adaptations to water stress (Saward, 1992; Spicer *et al*., 1993). Both in the tropical and paratropical belt, vegetation was possibly semi-arid or dry sub-humid, similar to open savanna forests with widely spaced coniferous trees, and a marked seasonal fluctuation in rainfall (Saward, 1992). Ancestral state reconstructions (Figure S7 and Appendix S1.6) suggest the conditions where the ancestor of the Core Caryophyllales inhabited were similar to modern dry sub-humid regions, with milder temperatures and high aridity indexes (0.57; see Appendix S1.5). The conditions are similar to the prevalent ones in the succulent biome worldwide, although the coincidence holds mostly for the New World, while in the Old World the estimated ancestral conditions mapped into modern climates reveal more a grassland type (Figure S7, also see Figure 15 in Schrire *et al*., 2005)).

Although adaptations to aridity were likely present early during the origin of the Core Caryophyllales, the succulent strategy appeared much later. We estimated that succulent lineages started diversifying at different time frames, from the Late Eocene to very recent dates such as the Pliocene (Appendix S1.3). However, the elevated variation both in confidence intervals for estimates as well as among estimates obtained using different approaches do not allow to confirm or reject the idea of a synchronous convergent diversification of succulents in response to an aridification trend (Arakaki *et al*., 2011).

Nevertheless, the results of our ENM analyses per succulent lineage (Figure 5) show there is not an equivalency amongst the climatic conditions present in the geographic regions of each succulent hotspot, suggesting local differences in the nature of the succulent syndrome and independent evolutionary trajectories for each lineage.

In all cases, our estimated dates for the diversification of succulents within the Caryophyllales are posterior to the estimated dates of diversification of legumes in the Succulent Biomes worldwide (Early Eocene, at 54.78 Ma Gagnon *et al*., 2019). These are suggested as grass-poor, fire intolerant dry tropical forests with an important presence of succulents (Schrire *et al*., 2005), although fire regime variables did not show informative in the determination of succulent records. Shrub and tree legumes have long been demonstrated to be fundamental for the lifecycle of succulents, particularly functioning as nurse plants (Valiente-Vanuet and Ezcurra, 1991); suggesting that once the legume framework was established, succulents started their profuse diversification.

Interestingly, although the diversification process of highly succulent lineages might have occurred at different time frames, our estimates for their origins (e.g. stem dates according to the estimates obtained with the more complete dataset: Anacampserotaceae 44.18 Mya, Cactaceae: 44.18 Mya, Didiereaceae: 42.92 Mya, see Appendix S1.3), as well as the origins of other succulent non-Caryophyllales (succulent *Euphorbia*, Euphorbiaceae: 39.59 Mya, (Bruyns *et al*., 2011); Agavaceae: 30-35 Mya (Good-Avila *et al*., 2006), succulent Asclepiadoideae at 30-35 Mya (Rapini *et al*., 2007), place their first appearance during the Eocene. The Eocene faced a cooling trend worldwide accompanied by a global drop off in atmospheric CO_2_ levels (DeConto and Pollard, 2003). These dates support the interesting hypothesis for the origin of the CAM as well as other carbon concentrating strategies, where early stem lineages that would give rise to highly succulent groups evolved as a strategy to deal with CO_2_ scarcity, and later colonized and diversified in semi-arid regions (Keeley and Rundel, 2003; Edwards and Ogburn, 2012).

### Succulent lineages might have diversified independently and by ecological fosters rather than aridity

Although we did not find a relationship with prevalence of succulence and diversification rates among Core Caryophyllales families, succulent lineages within them such as the Cactoideae (Cactaceae) or the Ruschioideae (Aizoaceae) present some of the highest diversification rates (see Appendix S1.2 and Magallón and Castillo, 2009; Hernández-Hernández *et al*., 2014; Valente *et al*., 2014). Accelerations in diversification rates have also been found in non-Caryophylales succulents such as *Agave* (Good-Avila *et al*., 2006) or *Euphorbia* (Horn *et al*., 2014). If as suggested here, succulent lineages were already preadapted to arid conditions, what can explain the relevance of the succulent syndrome to the diversification of these lineages? According to our estimated dates, succulent lineages diversified under climatic conditions that were already arid (Zachos *et al*., 2008). Several biotic or abiotic fosters of diversification have been proposed for succulents rather than aridity. For example, for Cactaceae it has been proposed that major diversification events were triggered by the evolution of novel growth forms and pollination syndromes (Hernández-Hernández *et al*., 2014). Pollination syndromes have also been proposed as a cause of the profuse diversification in *Agave* (Good-Avila *et al*., 2006; Flores-Abreu *et al*., 2019), and in the case of Aizoaceae, local edaphic adaptation to different soil types could have promoted the radiation of several succulent genera (Ellis *et al*., 2006; Ellis and Weis, 2006). More detailed studies of diverse succulent lineages are needed to understand their diversification fosters, with a particular focus on reproductive strategies.

Our ENM predictions and projections show that there are particular and distinctive climatic conditions on each succulent hotspot or high richness area per lineage, and they are non-equivalent, since they cannot inter-predict each other (Figure 5). Mean and median values for climatic variables for each succulent group differ considerably (Appendix S1.2). These differences, together with our ENM analyses results, show the relevance of an important discussion about the complex and unique set of climatic conditions existing under each of the geographic regions that we generalize as ‘deserts’. The abiotic variation among deserts is probably greater than for any other biome, largely because deserts are so widely spaced on the planet and have arisen for very different reasons (Ward, 2009). The high spatial and temporal variability of the abiotic environment in arid lands have important implications to desert life both at the ecological and evolutionary scales (Sandquist, 2014), and these gave rise to a tremendous plant diversity (i.e., life forms, functional diversity, reproductive and life strategies), and elevated diversification rates in several lineages.

In conclusion, our results show the versatility of the succulent plants, and reveal a little about the ecological complexity of this strategy. Although the whole Core Caryophyllales show a tendency towards inhabiting drylands, only few non-succulent species can withstand extremely arid conditions, as well as expand towards the more humid. The succulents’ climatic niche is contained within the non-succulents’, and is distributed more narrowly over the niche space, indicating a reduced environmental versatility possibly as a trade-off for increases in reproductive versatility. Although the ancestors of succulent CAM lineages might have originated during the Eocene, in response to a lower availability of CO_2_, the richest succulent lineages diversified later, possibly at different time frames, in different parts of the world under non-equivalent climatic conditions. All these results indicate that explanations of the evolution and diversification of succulents based solely on their convergence due to adaptation towards arid conditions might be oversimplistic. Succulent lineages are some of the richest and show the fastest diversification rates in the plant kingdom. Given their water and carbon balance, we consider our data support the idea that the succulent life strategy might provide advantages in terms of allowing these lineages broader capabilities to invest energy during the whole life cycle of the plant, for example, to explore more widely the reproductive niche space.

## Supplementary data

Supplementary data are available at JXB online.

**Extended Methods –** Details on estimation of dates analyses, classification of plants into different categories, statistical analyses, curation strategy

**Appendix S1 –** Databases & results (accession numbers, estimated dates, classifications, results of additional analyses)

**Appendix S2** – Aligned DNA matrices and phylogenetic trees

**Supplementary Figures –** Statistical analyses, traitgrams and ancestral reconstruction of climatic conditions map

**Supplementary Tables –** Distribution of species among categories of climate and succulence degree

## Acknowledgements

THH and MVC want to thank the support of LANGEBIO-UGA Director’s Office for support. MDA wants to thank CONACYT for graduated studies grant 487933.

## Author contributions

THH planned and designed the research, assembled databases, performed analyses, and wrote the manuscript, MN advised on statistical analyses, MD and MN performed statistical analyses, MVC and IL performed analyses.

